# Pupil responses reveal temporally distinct signatures of value updating and decision strategy

**DOI:** 10.64898/2026.06.13.732042

**Authors:** Shilpa Dang, Arezoo Pooresmaeili

## Abstract

Adaptive decision making depends on dopaminergic and noradrenergic systems supporting value learning and exploratory decision strategies, respectively, yet their contributions remain difficult to dissociate non-invasively. Here, we investigated whether pupil dynamics provide dissociable readouts of computational processes underlying value-based decision making across sensory modalities, potentially reflecting distinct neuromodulatory processes. Human participants performed a dynamic foraging task in which they chose between auditory, visual, or audio-visual options based on their estimated value, alongside a control task where choices were instructed. Behavior in the value-based task was well captured by a probabilistic choice model, revealing adaptive integration of reward history and a balance between exploration and exploitation. Pupil responses revealed temporally distinct computational signatures of decision strategy and value updating. Reaction time was associated with sustained pupil dilation throughout the decision process, whereas value differences between chosen and unchosen options selectively modulated pupil responses during stimulus evaluation and following feedback. These findings are consistent with computational processes linked to noradrenergic regulation of exploration–exploitation behavior and dopaminergic value updating, respectively. Both effects were significantly stronger during value-based than instructed decisions, indicating enhanced engagement of these computational processes when choices depended on learned reward values. Importantly, these effects were largely modality-independent, indicating a domain-general encoding of computational variables. Together, these findings identify pupil dynamics as a temporally sensitive and non-invasive marker of distinct computational stages underlying adaptive decision making and establish a framework for linking pupillometry to neuromodulatory theories of value learning and uncertainty processing.

## Introduction

Understanding how the brain dynamically balances value and uncertainty during decision making is a central question in cognitive neuroscience (Levallois et al., 2012; Padoa-Schioppa, 2011; Rangel et al., 2008). Value-based decisions require the integration of expected rewards, costs, and environmental structure, processes that are critically shaped by interacting neuromodulatory systems (Doya, 2008). In naturalistic settings, decision-relevant information is inherently multimodal and dynamic, requiring organisms to continuously update value estimates while adapting to changing environmental contingencies. In such environments, adaptive behavior depends on neural mechanisms that simultaneously evaluate reward value and uncertainty.

Two neuromodulatory systems have been proposed to play complementary roles in these computations. Dopaminergic signals encode expected reward and reward prediction errors (RPEs) (Schultz et al., 1997), particularly within regions such as the striatum (McClure et al., 2003; Montague et al., 1996), thereby motivating actions that maximize subjective value (Kawagoe et al., 1998). In parallel, the locus coeruleus–noradrenergic (LC–NE) system has been implicated in tracking uncertainty in action outcomes and facilitating adaptive exploitation-exploration strategies when action–outcome contingencies are volatile (Aston-Jones & Cohen, 2005; Doya, 2008). Consistent with its proposed role in regulating exploration–exploitation trade-offs, LC activity is more tightly coupled to behavioral responses than stimulus evaluation and correlates with trial-to-trial fluctuations in reaction time (RT), with shorter RTs often associated with exploitative and longer RTs with exploratory decision behaviors (Aston-Jones & Cohen, 2005; Rajkowski et al., 2004). Together, these systems provide a mechanistic framework for understanding how organisms balance reward maximization with behavioral flexibility under uncertainty.

While these neuromodulatory influences have been extensively characterized using invasive recordings, a growing body of work highlights the pupil as a non-invasive, temporally sensitive index of underlying neuromodulatory dynamics (Cremer et al., 2023; De Gee et al., 2014; Grujic et al., 2024; Reimer et al., 2016). Pupil diameter fluctuations have been primarily linked to activity in the locus coeruleus–noradrenergic (LC–NE) system and are increasingly recognized as reflecting both sensory-driven arousal during wakefulness and task-related behavioral states (Aston-Jones & Bloom, 1981; Bang et al., 2023; Manella et al., 2017). Consistent with adaptive gain theory (Aston-Jones & Cohen, 2005), pupil responses track LC-dependent regulation of task engagement and behavioral flexibility, including shifts between exploitative and exploratory behavioral states observed in both monkeys (Rajkowski et al., 1993) and humans (Gilzenrat et al., 2010). Additionally, recent studies suggest that pupil responses also capture signatures of dopaminergic computations underlying reward prediction error–guided learning (Van Slooten et al., 2018), reward–effort evaluation during decision making (Varazzani et al., 2015), and instrumental motivation for goal-directed actions (Grogan et al., 2020). This positions pupillometry as a promising tool for tracking latent computational variables and underlying neuromodulatory dynamics involved in value-based decisions.

Despite these advances, it remains unclear whether pupil dynamics primarily reflect general arousal or can dissociate computational processes associated with distinct stages of decision formation. In particular, the temporal evolution of pupil responses associated with value computations during choice, uncertainty-related decision processes, and outcome evaluation has not been clearly dissociated. Furthermore, although value-based decisions in natural environments often depend on information conveyed through multiple sensory modalities, it remains unclear whether pupil-linked computational signals are represented in a modality-general or modality-specific manner. This question is particularly relevant given evidence that both modality-specific and modality-independent representations have been observed in higher-order brain regions involved in decision making (Dang et al., 2024). Addressing these gaps is essential for linking computational processes underlying value-based decision making with physiological markers. Establishing such links may also enhance the translational and clinical utility of pupillometry, which has already demonstrated promising potential in studies involving clinical populations such as patients with Parkinson’s disease (Manohar & Husain, 2015; Tsitsi et al., 2023).

In the present study, we investigated how pupil responses track computational components of adaptive decision making in a multimodal environment. Participants performed a dynamic foraging task adapted from previous studies in which choices between auditory, visual, and audiovisual stimuli were guided by changing reward contingencies (Corrado et al., 2005; Dang et al., 2024; Serences, 2008). To dissociate value-related computations from more general task-evoked influences on pupil size, we additionally included a matched instruction-based control task in which participants selected explicitly instructed options rather than relying on subjective value estimates (**Fig. 1**).

**Figure 1.**
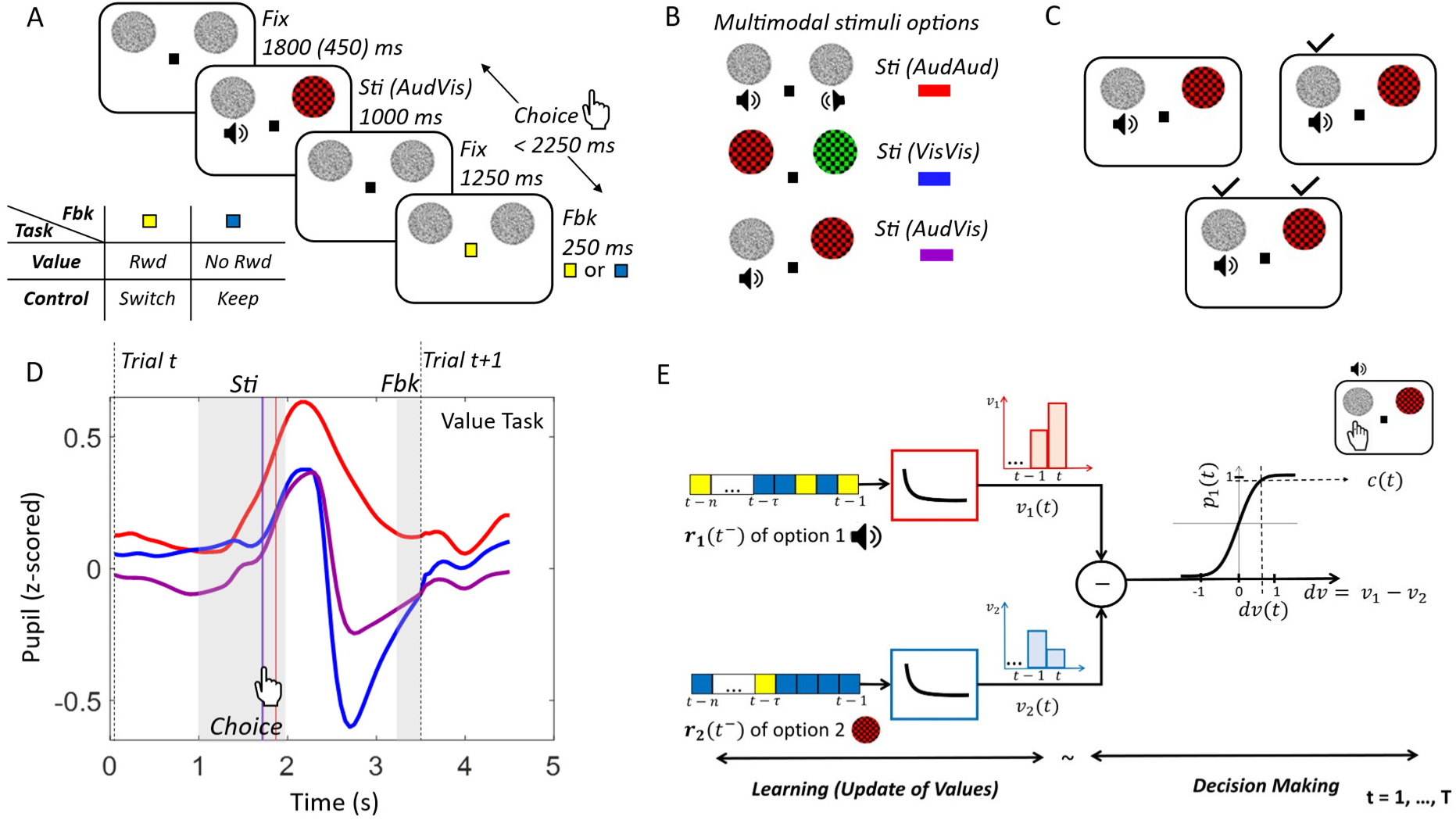
Experiment design and computational framework. **(A)** Schematic of an audio–visual (AudVis) trial in both behavioral tasks (value and control). Each trial began with a jittered fixation interval (Fix), followed by simultaneous presentation of stimulus options (Sti). Participants responded within a 2.25 s window, after which the fixation changed colour (yellow or blue) to convey feedback (Fbk). In the value task, yellow indicated reward (Rwd) and blue indicated no reward (No Rwd); in the control task, yellow and blue instructed participants to switch or keep their previous trial’s choice, respectively. Two white-noise placeholders were present throughout. **(B)** Examples of all stimulus modalities, where both options were auditory (AudAud), visual (VisVis), or one auditory and one visual (AudVis). When a visual option was shown, the corresponding placeholder was replaced by a coloured checkerboard. **(C)** In the value task, rewards were assigned independently and stochastically to each option (*S*_1_ or *S*_2_), such that a trial could contain rewards for none, one, or both options (illustrated by black check marks). On average, rewards occurred on 33% of trials and were distributed according to ratios of 3:1, 1:1, or 1:3. The control task followed an analogous structure, with switches assigned instead of rewards in an equiprobable manner with an average rate of 33%. **D)** Representative pupil response across a trial for all sensory modalities from a single participant in the value task. The grey shaded regions from 1–2 s and 3.25–3.5 s indicate the periods of stimuli options and feedback presentation, respectively. Responses are indicated by the hand icon. The solid red, blue, and purple vertical lines denote the mean reaction times for auditory, visual, and audio–visual trials, respectively. The dashed black vertical line denotes trial start. **E)** In the learning stage, reward histories of the two options (*r*_1_, *r*_2_) pass through identical exponentially decaying filters to compute their subjective values (*v*_1_, *v*_2_). At the choice stage, the difference between these values yields the differential value (*dv*), which determines the choice probability via a sigmoidal decision function (illustrated here for option 1), ultimately leading to the final choice.

We first quantified participants’ choice behavior using a probabilistic choice model that estimated subjective values from reward history and captured how participants balanced exploration and exploitation over time (Corrado et al., 2005). We then used these computational and behavioral signatures as predictors of pupil responses to examine how temporal pupil dynamics relate to distinct stages of decision making and whether these relationships differ across sensory modalities. Our hypothesis was that pupil dynamics would capture temporally dissociable signatures of value updating and uncertainty-related decision processes that generalize across sensory modalities, thereby providing a non-invasive window onto computational mechanisms associated with dopaminergic and noradrenergic function.

### Behavioral Results and Model Parameters

Participants performed a dynamic foraging task in which they repeatedly chose between two concurrently available stimulus options. The available options were drawn from two predefined stimulus sets (*S*_1_and *S*_2_) spanning auditory, visual, and audiovisual conditions (**Fig. 1, see Methods**). Reward availability was determined independently for each option according to a stochastic Poisson process, such that rewards were delivered on approximately one-third of trials. Across blocks, reward contingencies varied between the two options according to predefined reward ratios (1:3, 1:1, and 3:1), requiring participants to continuously track changing reward values and adapt their choices accordingly.

To characterize choice behavior, we quantified the proportion of choices allocated to *S*_1_relative to *S*_2_across reward contingencies and sensory modalities in the value task. Consistent with the matching law (Herrnstein & Baum, 1970), participants allocated their choices according to the relative reward rates of the two options, increasingly favouring *S*_1_ over *S*_2_ as *S*_1_: *S*_2_ reward ratio shifted from 1:3 to 1:1 to 3:1 (**Fig. 2A**)). A two-way repeated-measures ANOVA with factors reward ratio (1:3, 1:1, 3:1) and modality (Aud, Vis, AudVis) revealed a strong main effect of reward ratio on choice ratio (*F*[2,32] = 32.48, *p* = 1.51 x 10^-5^), with no significant interaction between reward ratio and modality (*F*[4,64] = 2.66, *p* = 0.11). Although there was a modest main effect of modality (*F*[2,32] = 4.89, *p* = 0.02), reflecting an overall shift in choice proportions in the auditory condition (Supplementary Information, **Table S1**), the influence of reward contingencies on choice behavior was comparable across modalities. We therefore collapsed behavioral data across sensory conditions in subsequent analyses (**Fig. 2B**).

**Figure 2.**
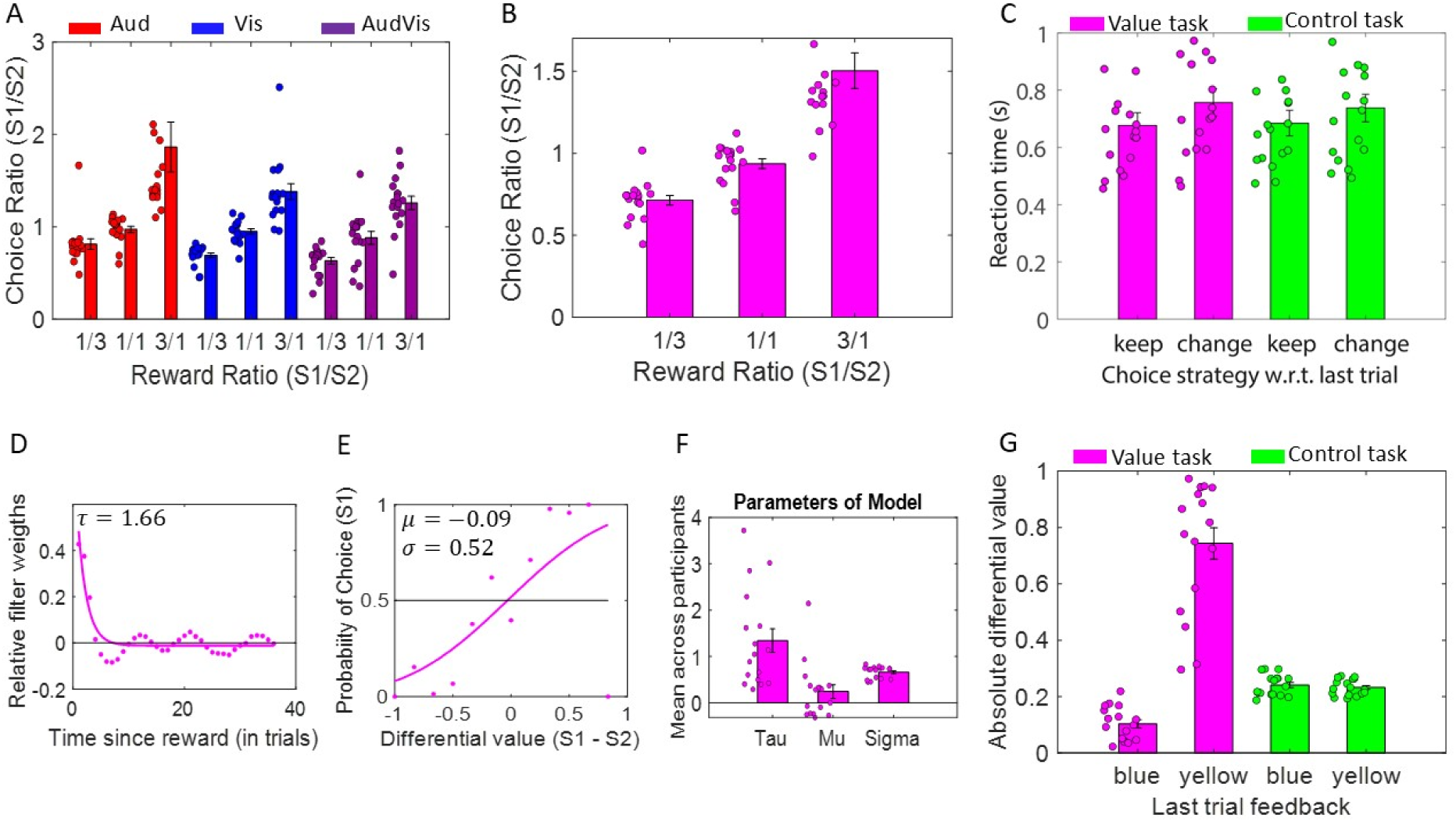
Behavioral results and model parameters. **(A)** Mean choice ratio across participants for each reward ratio (1:3, 1:1, 3:1) on stimuli options *S*_1_: *S*_2_, shown separately for each sensory modality in the value task (AudAud, VisVis, AudVis). Stimulus sets are defined as *S*_1_ = {low pitch, green, auditory} and *S*_2_= {high pitch, red, visual}. **(B)** Mean choice ratios from panel A collapsed across modalities, shown as a function of reward ratio in the value task. **(C)** Mean reaction times across participants for the value and control tasks, separated by whether participants repeated (“keep”) or switched (“change”) their choice relative to the previous trial, providing a clear behavioral signature of exploitative versus exploratory decision strategies. **(D)** Relative linear filter weights (dots) and their exponential fit (solid line) for a representative participant, illustrating how past rewards are weighted as a function of time in the value task. The parameter *τ* reflects the temporal decay timescale. **(E)** Relationship between differential value (*S*_1_ - *S*_2_) and the probability of choosing option *S*_1_ (dots), along with the sigmoidal fit (solid line) for the same participant. Parameters *µ* and *σ* capture choice bias and sensitivity to value differences, respectively. **(F)** Mean model parameters (*τ*, *µ*, *σ*) across participants. **(G)** Absolute differential value (*dv*) as a function of previous trial feedback (blue, yellow) for value and control tasks. In the value task, blue feedback indicated no reward and yellow feedback indicated reward, whereas in the control task, blue instructed participants to keep their previous choice and yellow instructed them to switch their previous choice. In panels **A**, **B**, **C**, **F**, and **G**, each dot represents an individual participant. Error bars indicate the standard error of the mean (s.e.m.) across participants.

For the control task, the analysis of choice behavior revealed that participants followed the task instructions, as choices were evenly distributed in accordance with the equal assignment of switches to the stimulus options, and showed no variation across sensory modalities (**Fig. S1**). A one-way repeated-measures ANOVA revealed no significant effect of modality on choice ratio (*F*[2,32] = 0.96, *p* = 0.39), indicating that participants’ choice pattern was comparable across auditory, visual, and audio-visual conditions. This suggests that modality did not influence decision behavior in instruction-based choices as well. For all further behavioral data analyses, choice data were collapsed across sensory modalities.

Reaction time (RT) data was analysed using a two-way repeated-measures ANOVA with factors task (value, control) and choice strategy (same vs. change relative to the previous trial), motivated by the link between exploit–explore dynamics and LC–NE system activity (Aston-Jones & Cohen, 2005). This analysis revealed a significant main effect of choice strategy (*F*[1,16] = 21.64, *p* = 2.66 x 10^-4^), with faster responses for repeated (keep) choices compared to switched (change) choices (**Fig. 2C**). In contrast, there was no significant main effect of task (*F*[1,16] = 0.08, *p* = 0.78) and no interaction between task and choice strategy (*F*[1,16] = 2.19, *p* = 0.16), indicating that this effect was consistent across both value-based and instruction-based decision contexts. Post-hoc comparisons confirmed that RTs were significantly shorter for keep versus change trials in both value (mean±s.e.m. = 0.68 s ± 0.04 s and mean±s.e.m. = 0.76 s ± 0.05 s, respectively, *p* = 3.19 × 10⁻⁴) and control (mean±s.e.m. = 0.68 s ± 0.04 s and mean±s.e.m. = 0.74 s ± 0.05 s, respectively, *p* = 5.71 × 10⁻³) tasks. Importantly, RTs did not differ between tasks within either strategy (keep: *p* = 0.72; change: *p* = 0.40). Together, these results suggest that decision strategy, rather than task context, primarily drives RT differences, consistent with a general mechanism underlying exploitative (faster) versus exploratory (slower) choices.

We next quantified the extent to which participants’ choices reflected dynamically updated reward values. To do so, we fitted a linear–nonlinear probabilistic choice model (referred to as the LNP model, see Methods) that inferred trial-by-trial subjective values (SVs) from reward history and predicted choice probabilities based on the difference in estimated SVs between the two available options (see **Fig. 1E** and *Methods*). For each participant, the model characterized reward learning through a temporal integration parameter (*τ*), which determined how strongly past rewards influenced current choices (**Fig. 2D**), and decision making through a sigmoidal choice function defined by a choice bias (*µ*) and sensitivity to value differences (*σ*; **Fig. 2E**). Across participants, the estimated timescale parameter *τ* (mean ± s.e.m. = 1.34 ± 0.25) was significantly greater than zero (*t*[16] = 5.27, *p* = 7.6 × 10⁻^5^), indicating that recent rewards had a stronger influence on choices than more distant past rewards (**Fig. 2F**). The bias parameter *µ* (0.24 ± 0.15) did not differ significantly from zero (*t*[16] = 1.67, *p* = 0.11), suggesting no systematic preference for either option. In contrast, the sensitivity parameter *σ* (0.66± 0.03) was significantly greater than zero (*t*[16] = 20.91, *p* = 4.8 × 10⁻^13^) and significantly different from one (*t*[16] = 10.77, *p* = 9.66 × 10⁻^9^), indicating that participants were sensitive to value differences and adopted a near-optimal balance between exploration and exploitation (with σ = 0 and σ ≫ 1 reflecting extreme exploitation and exploration strategies, respectively). Consistent with this strategy, participants harvested 93.00% (± 1.53% s.e.m.) of the available rewards. Importantly, model parameters were first estimated separately for each sensory modality within the value task and subsequently combined, yielding comparable results across modalities (see **Supplementary Fig. S2** and **Table S2**). Together, these findings demonstrate that participants’ choices were well explained by the LNP model, suggesting that behavior was guided by dynamically updated subjective value estimates derived from reward history across all conditions.

In the control task, participants followed the instructions conveyed by feedback with high accuracy (93.32% ± 1.94%, collapsed across keep and switch trials), indicating a clear understanding of the task rules. Because choices were explicitly instructed on a trial-by-trial basis, they were not expected to depend on past reward history, in contrast to the value task. This distinction was captured by the LNP model (**Fig. 2G**), which revealed a significant interaction between task type (value vs. control) and feedback type (blue vs. yellow) on updates in absolute differential values (*absDVs*; F[1,16] = 85.32, *p* = 8.19 x 10^-8^). In the value task, absDVs differed markedly as a function of feedback (mean ± s.e.m. = 0.11 ± 0.01 for blue feedback and 0.72 ± 0.05 for yellow feedback; *p* < 10⁻^8^), reflecting learning from reward outcomes. In contrast, no such difference was observed in the control task (0.24 ± 0.01 vs. 0.23 ± 0.01 for blue and yellow feedback, respectively; *p* = 0.15), consistent with the absence of value updating. These results demonstrate that the LNP model successfully dissociated feedback-driven learning in the value task from instruction-following behavior in the control task.

Overall, the behavioral results confirm that the value task elicited active, feedback-dependent learning of stimulus values across modalities, whereas the control task effectively minimized such processes, with participants relying solely on instructed choice rules.

### Dissociable pupil correlates of valuation and decision processes

We first examined whether pupil responses tracked trial-by-trial fluctuations in computational variables associated with value learning and decision strategy. To this end, we applied a time-resolved general linear model (GLM) to z-scored pupil responses (**Fig. 3A–C**). The GLM included parametric regressors capturing trial-by-trial fluctuations in subjective differential value (DV) and reaction time (RT), the regressors that were the main focus of our analyses and captured independent aspects of behavior (also see the **Supplementary Information**, under the analysis of collinearity). Additionally, unmodulated event-related regressor (EV) associated with common trial events such as stimulus presentation, motor response, and feedback delivery were also included (see **Fig. 1**). This approach allowed us to dissociate value-related, decision-related, and task-general arousal contributions to pupil dynamics across sensory modalities and tasks. Regression coefficients were estimated at each time point relative to trial onset and analysed at the group level to identify temporal intervals during which each regressor reliably modulated pupil responses (see ***Methods***).

**Figure 3.**
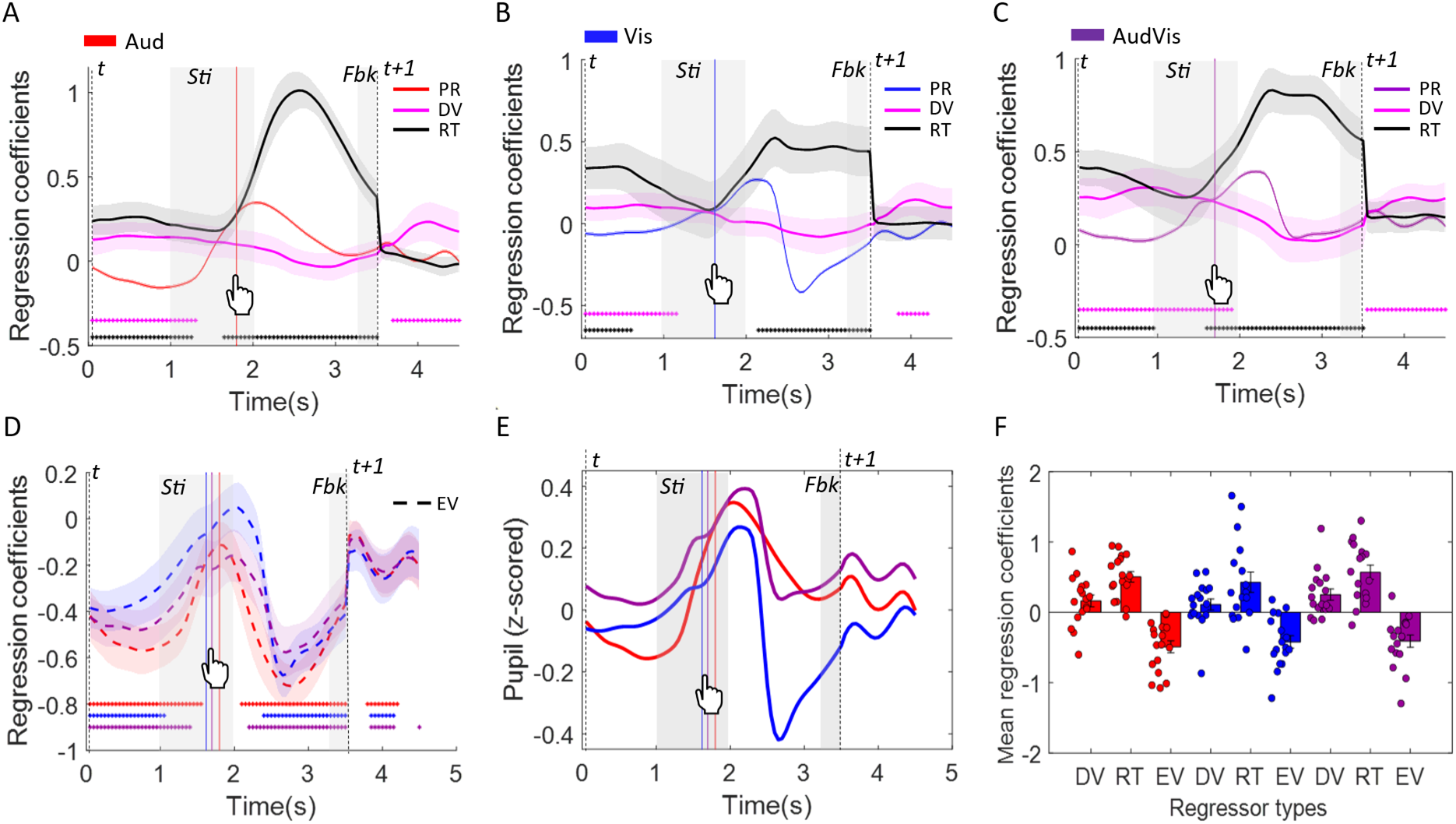
Pupil dynamics and regression estimates across sensory modalities in the value task. **(A–C)** Mean pupil response (z-scored) and time-resolved regression coefficients across participants during a trial for auditory (A; red), visual (B; blue), and audio-visual (C; purple) conditions. In each panel, traces represent the pupil response (PR; in modality-specific colour) and regression coefficients for the parametric regressors differential value (DV; magenta) and reaction time (RT; black), with shaded regions indicating ± s.e.m. across participants. Key trial events are indicated, including stimulus presentation (Sti; 1–2 s), response, and feedback (Fbk; 3.25–3.5 s). Vertical dashed lines mark trial onset and end. **(D)** Time-resolved regression coefficients for the unmodulated event (EV) regressor across modalities. The EV regressor captures task-general arousal responses linked to stimulus presentation, motor execution, and feedback delivery, thereby enabling isolation of value-and decision-related effects. Across panels **A–D,** horizontal significance bars indicate time points where coefficients differ significantly from zero (p < 0.05, corrected for multiple comparisons) at group-level. **(E)** Mean pupil responses (z-scored) shown in A-C across participants for auditory (red), visual (blue), and audio-visual (purple) trials are shown together to allow comparisons of PR across modalities. **(F)** Mean regression coefficients across participants, computed over trial time points showing significance at the group level, for all three regressors (DV, RT, and EV), shown separately for each modality (Auditory, Visual, Audio–visual). Each dot represents an individual participant; error bars indicate the standard error of the mean (s.e.m.) across participants.

After accounting for task-evoked arousal (EV, **Fig. 3D**), we found that both reaction time (RT) and subjective value difference (DV) significantly modulated pupil dynamics over distinct temporal intervals within a trial of the value task (**Fig. 3A–C**). Across auditory, visual, and audiovisual conditions, RT exerted a robust and sustained influence on pupil responses, with significant effects observed both before the behavioral response (∼0–1 s) and during the post-decision period (∼2–3.5 s; *p* < 0.05, corrected for multiple comparisons; **Fig. 3A–C**).

Notably, significant RT-related modulation emerged before the behavioral response (∼0–1 s), indicating that pupil dynamics were sensitive to processes unfolding during decision formation. Given the established relationship between RT and exploration–exploitation behavior, this early effect may reflect trial-by-trial variation in pre-decisional processes that influence how rapidly a choice is reached. RT-related modulation persisted during the post-decision period (∼2–3.5 s; *p* < 0.05, corrected), suggesting that pupil dynamics continued to track processes associated with the completed decision. One possibility is that this sustained effect reflects uncertainty-related processes that remain active while participants await feedback regarding the outcome of their choice. Consistent with this interpretation, the RT effect decreased sharply following feedback presentation, suggesting that the processes linked to RT were largely resolved once outcome information became available.

In contrast, DV-related effects (*p* < 0.05, corrected) showed a temporally selective profile, emerging early in the trial and persisting through the initial phase of stimulus processing (∼0–1.5 s), before re-emerging during the post-feedback period (∼3.5 s onwards). This temporal pattern is consistent with the role of subjective value in both choice formation and feedback-driven learning. The early effect likely reflects value-related computations during stimulus evaluation, whereas the late effect coincided with the period immediately following feedback, when stimulus–value associations are updated based on outcome information (Dang et al., 2024). Although smaller in magnitude than RT-related effects, DV-related modulation was reliably observed across all sensory modalities. Together, the distinct temporal profiles of RT and DV effects suggest that pupil dynamics track separable computational components of value-based decision making, including decision strategy and value updating.

Despite minor differences in the precise temporal profiles across auditory, visual, and audio-visual conditions in the value task, the overall pattern of results was qualitatively similar across modalities, indicating a largely modality-independent encoding of the task-related variables in pupil responses (**Fig. 3A–D**). To quantify this, we averaged regression coefficients within the significant time windows identified for each regressor and performed a two-way repeated-measures ANOVA with factors modality (auditory, visual, audiovisual) and regressor type (DV, RT, EV) (**Fig. 3F**). This analysis revealed a significant main effect of regressor type (F[2,32] = 40.56, p = 1.68 x 10^-9^) and a significant main effect of modality (F[2,32] = 7.60, p = 0.002), with no interaction between the two factors (F[4,64] = 0.20, p = 0.94). Post-hoc comparisons showed that both DV-related and RT-related coefficients were significantly larger than EV coefficients across all modalities (all p < 0.005), indicating that value-related and decision-related signals contributed more strongly to pupil dynamics than task-general event-driven arousal effects (**Table S3**). RT effects were generally larger than DV effects, although this difference reached significance only in the auditory condition. Critically, no significant differences were observed across modalities within each regressor type (all p ≥ 0.86), indicating that the encoding of DV, RT, and EV signals in pupil responses was largely modality-independent in the value task.

These results complement the time-resolved analyses (**Fig. 3A–D**), where RT-related effects dominated both pre- and post-decisional trial periods, reflecting decision formation and uncertainty processes, while DV-related effects emerged around stimulus evaluation and re-emerged following feedback, consistent with ongoing value updating. In contrast, EV-related activity was relatively weaker, indicating baseline arousal associated with task events. Together, these findings demonstrate that pupil dynamics are primarily driven by computational variables underlying value-based decision making, with a robust and modality-general encoding of both feedback-related value updating and decision strategy signals.

### Cross-task analysis

Next, we examined whether pupil-linked signatures of value updating (DV; putatively linked to dopaminergic signals), decision strategy (RT; putatively LC–NE mediated), and general event-related arousal (EV) differed between value-based and instruction-based decision contexts, thereby allowing us to isolate processes specific to value-based learning. Examining the temporal evolution of regression coefficients (collapsed across sensory modalities; **Fig. 4A–C**) revealed a clear and dissociable unfolding of signals across the trial, with important differences between tasks.

**Figure 4.**
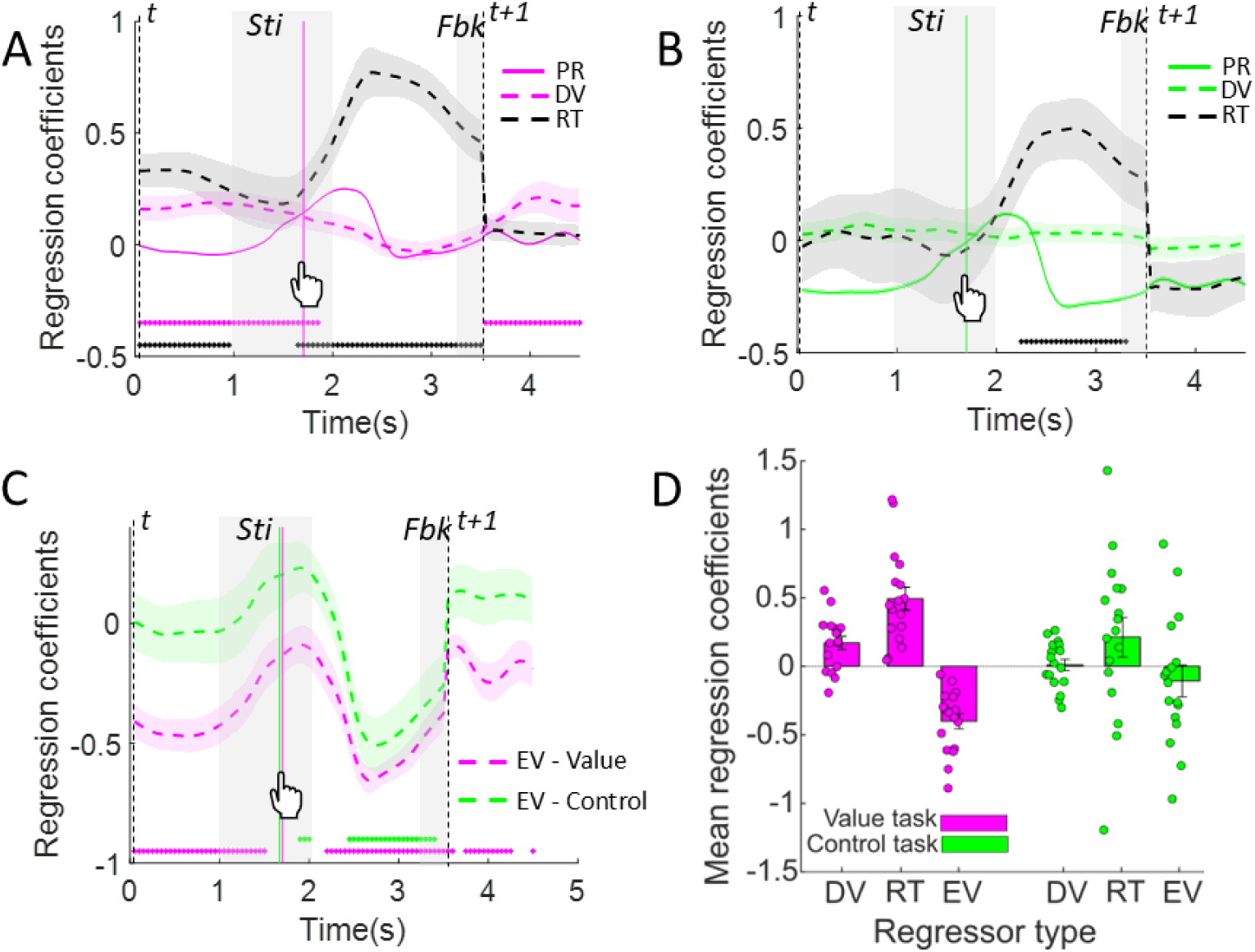
Pupil dynamics and regression estimates across value and control tasks. **(A–B)** Mean pupil response (z-scored) and time-resolved regression coefficients across participants during a trial for the value task (A; magenta) and control task (B; green). In each panel, traces represent the pupil response (PR; task-specific colour) and regression coefficients for the parametric regressors differential value (DV; task-specific colour dashed line) and reaction time (RT; black dashed line), with shaded regions indicating ± s.e.m. across participants. Key trial events are indicated, including stimulus presentation (Sti; 1–2 s), response, and feedback (Fbk; 3.25–3.5 s). Vertical dashed lines mark trial onset and end. **C)** Time-resolved regression coefficients for the unmodulated event (EV) regressor for value (magenta dashed) and control (green dashed) tasks. The EV regressor captures task-general arousal responses linked to stimulus presentation, motor execution, and feedback delivery, thereby enabling isolation of value- and decision-related effects. Across panels **A–C**, horizontal significance bars indicate time points where coefficients differ significantly from zero (p < 0.05, corrected for multiple comparisons) at the group level. **D)** Mean regression coefficients across participants, computed over trial time points showing significance at the group level, for all three regressors (DV, RT, and EV), shown separately for value (magenta) and control (green) tasks. Each dot represents an individual participant; error bars indicate the standard error of the mean (s.e.m.) across participants.

RT-related effects exhibited a strong and sustained modulation during the response window (∼2–3.25 s; p < 0.05, corrected for multiple comparisons, see *Methods*) in both tasks, peaking during post-decision period and thereafter gradually declining, consistent with post-decisional uncertainty. Importantly, an early RT significance (∼0–1 s; p < 0.05, corrected for multiple comparisons) appeared only in the value task, not in the control task, which suggested that it was specifically tied to the fluctuating dynamics of value-guided decision formation rather than trivial instruction-based decision formation. In a dynamic reward environment, participants continuously evaluate options based on recent reward history, and this ongoing valuation process influences both decision speed (RT) and arousal (pupil size) from the very beginning of the trial. Thus, the pupil signal suggests an anticipatory coupling between valuation and action. In contrast, in the control task, where decisions were externally instructed and did not depend on internal value computations, this early coupling between RT and pupil was absent. This reinforces the interpretation that the early RT effect in the value task reflected endogenous/internal decision dynamics, likely linked to uncertainty and exploit–explore state, rather than passive instruction execution. Overall, the early RT significance suggests that pupil dynamics are not just reactive but predictive, indexing the internal decision state that will ultimately determine how quickly a choice is made.

DV-related effects, in contrast, showed a temporally structured and task-dependent profile (p < 0.05, corrected for multiple comparisons). In the value task, DV modulated pupil responses during early stimulus evaluation (∼0–2 s) and re-emerged following feedback (∼3.5 s onwards), consistent with ongoing value computation and feedback-driven updating of stimulus–value associations. This late component reflected the updating of subjective value that carries forward into the next trial. In the control task, however, DV-related modulation was hardly ever significant in any time window, consistent with the absence of value-based learning (**Fig. 4B**). Finally, EV-related responses exhibited a transient, event-locked profile aligned with stimulus and feedback periods, with a comparable profile across tasks.

Together, these temporal patterns indicate that pupil dynamics are dominated by a robust, task-general decision-related signal (RT), while selectively revealing task-specific signatures of value-guided decision formation. The latter was evident in the early anticipatory RT effect and feedback-linked DV modulation that are preferentially expressed in the value task, alongside more general event-driven arousal responses.

To formally test these effects, we performed a two-way repeated-measures ANOVA on mean regression coefficients (averaged across significant time windows with respect to the value task) with factors task (value, control) and regressor type (DV, RT, EV) (**Fig. 4D**). This analysis revealed a significant main effect of regressor type (F[2,32] = 13.16, p = 6.77 x 10^-5^), no significant main effect of task (F[1,16] = 2.42, p = 0.14) and a significant interaction between task and regressor type (F[2,32] = 6.08, p = 0.006), indicating that the relative contribution of different regressors to pupil dynamics differed across tasks.

Post-hoc comparisons further clarified the nature of the task × regressor interaction (Table S4). Within the control task, pairwise comparisons between regressors (DV, RT, and EV) revealed no significant differences, indicating a relatively undifferentiated contribution of value-, decision-, and event-related factors to pupil responses. We next compared the strength of each regressor across tasks. DV coefficients were significantly larger in the value task than in the control task (p = 0.04), consistent with the engagement of value-based computations during reward-guided decisions. RT coefficients were also greater in the value task (p = 0.02), indicating stronger decision-related modulation of pupil responses when choices depended on learned reward values. In contrast, EV coefficients were more negative in the value task (p = 0.03), suggesting that value- and decision-related factors accounted for a greater proportion of pupil variance during reward-guided behavior. Together, these findings indicate that pupil dynamics in the value task were more strongly shaped by computational variables associated with learning and decision formation than by task-evoked responses alone.

Together, these findings indicate that pupil dynamics reflect both task-general decision processes and task-specific value computations. While reaction-time-related signals constituted the strongest contributor to pupil fluctuations across tasks, value-related signals were selectively amplified during reward-guided decision making, supporting a dissociation between general decision dynamics and feedback-driven valuation processes.

## Discussion

The present study set out to determine whether pupil dynamics can dissociate and track core computational components of value-based decision making, namely feedback-driven value updating and ensuing decision strategy, within a unified framework and in a multimodal environment. By combining computational modelling with time-resolved pupillometry, we demonstrate that pupil responses carry temporally and functionally dissociable signatures of these processes. Specifically, reaction time, indexing decision strategy and uncertainty, exerted a robust and sustained influence on pupil dilation during early decision formation window and around choice execution, whereas differential value (DV), indexing feedback-related valuation, showed temporally specific modulation during early stimulus evaluation and following feedback. Importantly, these effects were largely modality-independent and exhibited clear task-dependent differences. Computational signatures were more strongly differentiated during value-based decisions, whereas instruction-based decisions showed comparatively undifferentiated contributions of value-, decision-, and event-related processes. Together, these findings suggest that pupil dynamics provide a temporally resolved readout of the computations underlying adaptive decision making and support the use of pupillometry as a non-invasive tool for studying neuromodulatory processes associated with value learning and uncertainty.

Our findings align with influential accounts of the Adaptive Gain Theory, which posit that the locus coeruleus–noradrenergic (LC–NE) system regulates the balance between exploitation and exploration (Aston-Jones & Cohen, 2005). The strong and sustained modulation of pupil responses by RT, particularly during the pre-decisional period, is consistent with evidence linking pupil dynamics to uncertainty-related processes and fluctuations in behavioral state. Longer RTs, associated with exploratory or uncertain decisions, were accompanied by larger pupil dilations, in line with prior work linking LC activity to variability in response latency (Aston-Jones & Cohen, 2005; Rajkowski et al., 2004). Notably, the presence of early RT-related modulation in the value task, but not in the control task, suggests that pupil dynamics capture anticipatory decision states arising from internal valuation processes rather than purely motor preparation. This extends previous observations by demonstrating that LC-linked signals can reflect pre-decisional computations in dynamic environments. Furthermore, although RT-related effects were present in both tasks, they were significantly greater in the value task, indicating enhanced engagement of decision-related processes when choices depended on learned reward values.

In contrast, DV-related effects exhibited a temporally structured profile, emerging during early stimulus processing and reappearing following feedback. This pattern is consistent with the role of dopaminergic systems in encoding expected value and reward prediction errors (Montague et al., 1996; Schultz et al., 1997). Although pupil diameter is more directly linked to LC–NE activity, accumulating evidence suggests that it can indirectly reflect dopaminergic influences, particularly through interactions between neuromodulatory systems (Van Slooten et al., 2018; Varazzani et al., 2015). The late DV-related pupil modulation observed after feedback likely indexes the updating of stimulus–value associations, a key component of reinforcement learning. This interpretation is further supported by the absence of comparable DV effects in the control task, where value updating was not required. Moreover, the lack of significant differentiation among DV-, RT-, and EV-related coefficients in the control task indicates that pupil dynamics did not preferentially encode any specific computational component when decisions were guided solely by instructions. In contrast, the value task exhibited clear separation among these components, suggesting that computational specialization of pupil responses emerges primarily in the context of reinforcement-driven learning. Thus, our results suggest that pupil dynamics capture not only decision-related uncertainty but also feedback-driven learning signals.

A central contribution of this study is the demonstration that these computational signals are largely invariant across sensory modalities. Despite differences in auditory, visual, and audio-visual inputs, both RT- and DV-related pupil effects showed similar temporal profiles and magnitudes. This modality-independent encoding suggests that the processes reflected in pupil dynamics operate at the level of abstract decision variables rather than sensory-specific representations, consistent with evidence that pupil responses track task contingencies and reward-guided behavior independent of stimulus modality (Antono et al., 2022). This view is consistent with prior work showing that LC activity reflects global brain states and task utility rather than modality-specific processing (Aston-Jones & Bloom, 1981; Reimer et al., 2016). By extending these findings to a dynamic foraging paradigm, we provide evidence that pupil-linked neuromodulatory signals support flexible decision making across complex, multimodal environments.

Finally, these findings have broader implications for the use of pupillometry in clinical and translational settings. Alterations in dopaminergic and noradrenergic function are implicated in disorders such as Parkinson’s disease, where deficits in reward processing and decision making are prominent (Manohar & Husain, 2015). By linking pupil dynamics to computational markers of valuation and decision strategy, the present work contributes to a growing framework for non-invasive assessment of neuromodulatory dysfunction. Future studies could extend this approach to patient populations and more naturalistic decision contexts.

In summary, our results demonstrate that pupil dynamics encode dissociable, temporally structured, and modality-general signatures of value updating and decision strategy. Crucially, such computational dissociation emerged primarily during reward-guided learning, whereas instruction-guided choices lacked clear differentiation among value-, decision-, and event-related components. These findings bridge computational models of behavior with physiological markers of neuromodulation, establishing pupillometry as a powerful tool for studying the mechanisms underlying adaptive decision making in complex environments.

## Methods

### Participants

Twenty-one healthy subjects (7 male and 14 female, age 19 to 45 years; mean ± SD age = 25.10 ± 3.79 years) participated in the eye-tracking experiment for financial compensation up to a maximum of 22€ based on their behavioral performance in the value-based decision-making task (value task). All were right-handed and had normal or corrected-to-normal vision and were naïve to the hypothesis of the project. Before the experiment started and after all procedures were explained, participants gave a written informed consent and completed a practice session. The study was performed at the European Neuroscience Institute, Göttingen (ENI-G). All procedures were approved by the local ethics committee of the *University Medical Center Göttingen*, under the proposal number 15/7/15.

Four participants were excluded resulting in data from 17 subjects included in our final sample: one participant had poor accuracy (45.77%) in control task and for three participants more than 10% of trials had to be removed due to poor fixation.

### Experimental Design and Procedures

The study comprised two decision-making paradigms: a value-based task (value task) and an instruction-based task (control task), administered across two sessions on the same day (**Fig. 1 A–C**). The design of the experiments, procedures, and the computational models of behavior were largely identical to a previous fMRI study, and are detailed in (Dang et al., 2024), with a brief explanation below.

Each session included 12 blocks of 72 trials. Nine blocks were allocated to the value task (one block per reward ratio: {1:3, 1:1, 3:1} per sensory modality), followed by 3 blocks of the control task. In both tasks, participants made binary choices between stimuli presented within or across sensory modalities (in separate blocks and pseudorandomized): auditory–auditory (*AudAud*), visual–visual (*VisVis*), and audio–visual (*AudVis*).

The choice set comprised two auditory and two visual stimuli (**Fig. 1B**). Auditory stimuli included a low-pitch (LP) sawtooth tone (294 Hz) and a high-pitch (HP) sinusoidal tone (1000 Hz), delivered through an over-ear headphone (HAD 280 audiometry headphones, Sennheiser). Visual stimuli consisted of two contrast-reversing checkerboards (green–black and red–black), flickering at 8 Hz (Serences, 2008), presented within circular apertures of 4° radius, and at 10° horizontal eccentricity and 5° above the center of the screen.

The value and control tasks were matched in all respects except for the interpretation of the feedback signal (**Fig. 1A**). In both tasks, a trial started with a fixation period (mean 1.8 ± 0.45 s), followed by stimulus presentation (1 s), a response window (2.25 s), and feedback (0.25 s). On each trial, two stimulus options were presented simultaneously on either side of fixation, and participants indicated their choice using the left or right arrow keys of a keyboard. The spatial position of the options was pseudo-randomised across trials. In the value task, feedback indicated whether the chosen option was rewarded (yellow) or unrewarded (blue). In the control task, feedback instructed behavior on the subsequent trial rather than conveying reward information: yellow indicated that participants should switch their previous choice, whereas blue indicated that they should repeat it. The first trial of each control block required a random choice.

To create a dynamic reward environment, rewards were assigned independently to each option according to a Poisson process (Corrado et al., 2005) (**Fig. 1C**). Within each block, rewards were distributed between the two options according to one of three reward ratios (1:3, 1:1, or 3:1), which were pseudo-randomised across sensory modality conditions.

To encourage adaptive learning and exploration, the task incorporated both a baiting procedure and a change-over delay (COD) (Corrado et al., 2005; Serences, 2008). Under the baiting procedure, rewards assigned to an option remained available until collected, discouraging persistent selection of the currently more rewarding option. The COD imposed a one-trial delay on reward delivery following a switch between options, discouraging simple alternation strategies. Because participants were aware that rewards could not be obtained immediately after a switch, trials following a switch were excluded from subsequent analyses in the value task.

At the end of each block, participants were shown their accumulated reward, with each rewarded trial represented as a yellow square worth 5 cents. After completing the experiment, participants received their total earnings, which were up to a maximum of €22 (€11 per session).

Switch instructions in the control task were assigned independently and stochastically to the options with equal probability, resulting in an average switch rate of 33%. Unlike the value task, no baiting or change-over delay mechanisms were applied. At the end of each block, participants were shown their performance, reflecting how accurately they followed the instructed choices.

### Computational Model of Choice

Choice behavior in the value task was quantified using a linear–nonlinear–probabilistic (LNP) model adapted from previous studies (Corrado et al., 2005; Herrnstein & Baum, 1970; Serences, 2008), and detailed in (Dang et al., 2024). The model estimated trial-by-trial subjective values for the two stimulus options from their recent reward history using a temporal weighting function, such that more recent rewards contributed more strongly to current value estimates (**Fig.1D**). The temporal profile of reward integration was characterised by a decay time constant (*τ*), which reflects the timescale over which past rewards influence current choices.

For each trial, the model computed a differential value,

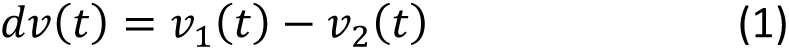

where (*v*_1_) and (*v*_2_) denote the estimated subjective values of the two available options. This differential value served as the decision variable and was mapped onto choice probability using a sigmoidal choice function (shown for an example participant in **Fig. 1E**). The resulting fit yielded two additional parameters: (*µ*), reflecting a systematic choice bias, and (*σ*), reflecting sensitivity to value differences. Lower values of (*σ*) indicate more deterministic, exploitative choices, whereas higher values indicate greater exploratory behavior.

To examine whether participants adopted such strategies, we analysed the fitted parameters *τ*, *µ* and *σ* with respect to behavioral data (**Fig. 2D–F**). The same LNP model was also fitted to the control task choices and compared absolute differential values (*absDVs*) across value and control tasks as a function of previous-trial feedback (**Fig. 2G**). Absolute DVs were used as a symmetric measure of subjective preference across options and were subsequently used as predictors in the pupillometry analyses.

### Pupil data acquisition and preprocessing

Pupil data (diameter) from the right eye was recorded at 1000 Hz using an EyeLink 1000 Tower Mount (SR Research, Ontario, Canada). Data from each participant was collected in two identical sessions in a single day with a 15-min break between the two sessions. Each session consisted of four runs (first three runs: 9 blocks of value task, last run: 3 blocks of control task). The eye-tracker was calibrated at the start of each run (9-point calibration). Saccades and blinks were detected using the standard EyeLink software and custom routines for preprocessing eye-tracking data in MATLAB (version 9.9; The Math Works, Natick, MA, USA). Linear interpolation was used to remove durations of signal loss due to blinks using interpolation window of half-width 100ms pre- and post-blink. The interpolated signal was low pass filtered at cutoff frequency 5 Hz using third-order Butterworth filter, z-scored per participant, and resampled to 20 Hz. Further, to ensure that analysis was not affected by eye movements, those trials were not included in which participants made a saccadic eye movement away from the fixation point more than 3.3 ° (percentage removed trials, mean ± SD = 5.16% ± 5.42%) (Palaskar and Dang, 2024; Van Slooten et al., 2018). Participants for whom more than 10 % of trials were lost were removed from further analysis (N=3).

### Pupil regression analysis

To examine how pupil dynamics track trial-by-trial fluctuations in computational variables underlying value-based decision making, we employed a time-resolved general linear model (GLM) framework. Our primary aim was to dissociate the contributions of feedback-related value updating and ensuing decision processes to pupil responses. Specifically, we quantified value-related signals using the trial-wise subjective value difference between the chosen and unchosen options (DV), derived from the LNP model, and decision-related processes using reaction time (RT), putatively reflecting exploitative versus exploratory choice dynamics.

For each participant, GLM analysis was performed on preprocessed, z-scored pupil time series, combining data from both value and control tasks. At each time point within a trial, pupil responses were modeled using three regressors: (i) an unmodulated event EV regressor capturing stimulus-driven and task-evoked arousal common to all trial events, (ii) a parametric DV regressor indexing trial-by-trial fluctuations in subjective value difference, and (iii) a parametric RT regressor indexing trial-by-trial variability in decision latency.

The EV regressor was included to account for unmodulated, event-related pupil responses associated with stimulus presentation, motor response, and feedback presentation, thereby isolating task-general arousal components. The EV regressor was modelled as 1 during events and 0 otherwise. The DV regressor captured value-dependent modulation of pupil size, based on the difference between the estimated subjective values of the chosen and unchosen options on each trial, reflecting underlying dopaminergic dynamics. These value estimates were obtained from the LNP model and z-scored within participants. The RT regressor captured variability in decision-related processes, where shorter RTs are often associated with exploitative choices and longer RTs with exploratory decisions, putatively reflecting underlying locus coeruleus–noradrenergic (LC–NE) dynamics. Both parametric regressors were mean-centered and entered simultaneously into the model to dissociate their independent contributions.

The GLM modeled each sensory modality (auditory, visual, and audio-visual) separately, such that regressors for a given modality were assigned non-zero values only for trials belonging to that modality and zero otherwise. This allowed us to estimate modality-specific regression coefficients while controlling for shared variance across conditions. Regression coefficients (beta weights) were estimated at each time point relative to trial onset, yielding time-resolved measures of the contribution of each regressor to pupil dynamics.

For group-level inference, we employed a random-effects analysis using a summary-statistic approach, performing one-sample t-tests against zero at each time point across participants (Penny & Holmes, 2003). To correct for multiple comparisons across time, we employed a non-parametric cluster-based permutation test. Time points exceeding an initial threshold (p < 0.05, uncorrected) were grouped into contiguous clusters. Cluster-level statistics were computed as the sum of t-values within each cluster and compared against a null distribution generated via permutation (1000 iterations). Clusters were considered significant if their cluster-level statistic exceeded the 95th percentile of the null distribution (p < 0.05, corrected for multiple comparisons). Significant time intervals were defined as those surviving this correction and were used to characterise when DV-, RT-, and EV-related signals reliably modulated pupil responses in both value and control tasks (**Fig. 3-4**).

### ANOVA on Mean Regression Coefficients

To formally quantify contributions of computational and event-related signals to pupil dynamics, we conducted additional ANOVA-based analyses on regression coefficients derived from the time-resolved GLM.

To compare the relative contributions of parametric and event-related components to pupil dynamics across modalities in the value task, we conducted an additional analysis on the regression coefficients obtained from the time-resolved GLM (**Fig. 3**). For each participant, regression coefficients corresponding to the three regressors—trial-by-trial differential value, trial-by-trial reaction time, and the unmodulated event regressor—were averaged across time points that showed significant effects at the group level (see Pupil regression analysis). These mean coefficients were computed separately for each sensory modality (auditory, visual, and audio-visual) and entered into a two-way repeated-measures ANOVA with within-subject factors modality (auditory, visual, audio-visual) and regressor type (DV, RT, EV). Post-hoc pairwise comparisons were performed using Bonferroni correction to examine differences between regressors within each modality and across modalities within each regressor type. This analysis allowed us to assess whether the strength of parametric (DV, RT) and event-driven (EV) contributions to pupil responses differed across modalities, thereby testing for modality-specific versus modality-general encoding of task-relevant variables in the value task.

Next, to dissociate pupil-linked neuromodulatory signatures associated with reward-guided valuation from those reflecting general task-related perceptual/motor and decision processes, we performed a cross-task analysis comparing the value and instruction-based (control) tasks using a unified framework (**Fig. 4**). Given the absence of modality-specific interactions in prior analyses, regression coefficients for each participant (corresponding to DV, RT, and EV regressors) were first collapsed across sensory modalities individually for both tasks. Thereafter, these coefficients were averaged within group-level significant time windows of the value task (identified from the time-resolved analysis) to obtain a single summary measure per participant, regressor, and task. The resulting mean regression coefficients were entered into a two-way repeated-measures ANOVA with factors task (value, control) and regressor type (DV, RT, EV). This analysis tested whether pupil-linked signatures of value updating (DV; putatively dopaminergic), decision strategy (RT; putatively LC–NE mediated), and general event-related arousal differed between reward-guided and instruction-driven decision contexts, thereby isolating neuromodulatory signals specific to value-based learning. Post-hoc comparisons (Bonferroni-corrected) were conducted to further characterize differences between regressors within each task and between tasks for each regressor.

## Acknowledgements

We are grateful to Igor Kagan for helpful discussions on the experimental design. This work was conducted at the European Neuroscience Institute Göttingen, Germany, and was supported by an ERC Starting Grant (No. 716846) awarded to AP.

## Authors’ contributions

SD and AP conceptualized the project and designed the task. SD conducted the experiments, preprocessed and analyzed the data. SD and AP interpreted the results and wrote the first draft of the manuscript. Both authors revised the manuscript. AP acquired funding.

## Supplementary Material

### Supplementary Text and Tables

#### Choice ratios across sensory modalities of the value task

Results shown in **Fig.2A**, indicate a strong effect of reward ratio across all sensory modalities of the value task on the choice ratio of *S*_1_ relative to *S*_2_. As indicated by an ANOVA analysis, there was a significant effect of reward ratio and no interaction with the sensory modality. However, a weaker but statistically significant effect of modality was also found. Inspection of post-hoc pairwise comparisons (**Table S1**) shows that this effect was mainly driven by a subtle bias toward choosing option *S*_1_in auditory modality (low-pitch tone). This tendency was not consistently significant across reward ratios but was more pronounced at the extreme ratios (1:3 and 3:1) of the value task.

**Table S1.** Post-hoc comparisons.

| Table S1. Post-hoc comparisons |  |  |  |  |
| --- | --- | --- | --- | --- |
| Factor<br>(Modality or<br>Reward ratio) | Comparison between<br>levels | Difference | Standard error | p-value |
| Auditory | 1:3 – 1:1 | -0.16 | 0.08 | 0.18 |
| Auditory | 1:1 – 3:1 | -0.87 | 0.30 | 0.03 |
| Auditory | 1:3 – 3:1 | -1.03 | 0.23 | $1.41 \times 10^{-3}$ |
| Visual | 1:3 – 1:1 | -0.25 | 0.03 | $1.09 \times 10^{-6}$ |
| Visual | 1:1 – 3:1 | -0.42 | 0.09 | $7.37 \times 10^{-4}$ |
| Visual | 1:3 – 3:1 | -0.67 | 0.10 | $1.43 \times 10^{-5}$ |
| Audio-Visual | 1:3 – 1:1 | -0.23 | 0.06 | $5.00 \times 10^{-3}$ |
| Audio-Visual | 1:1 – 3:1 | -0.39 | 0.06 | $2.11 \times 10^{-5}$ |
| Audio-Visual | 1:3 – 3:1 | -0.62 | 0.07 | $2.39 \times 10^{-7}$ |
| 1:3 | Auditory – Visual | 0.11 | 0.06 | 0.21 |
| 1:3 | Visual – Audio-Visual | 0.05 | 0.03 | 0.29 |
| 1:3 | Auditory - Audio-Visual | 0.17 | 0.06 | 0.05 |
| 1:1 | Auditory – Visual | 0.02 | 0.04 | 1.00 |
| 1:1 | Visual – Audio-Visual | 0.08 | 0.08 | 1.00 |
| 1:1 | Auditory - Audio-Visual | 0.17 | 0.06 | 0.05 |
| 3:1 | Auditory – Visual | 0.47 | 0.27 | 0.31 |
| 3:1 | Visual – Audio-Visual | 0.11 | 0.11 | 1.00 |
| **3:1** | Auditory - Audio-Visual | 0.58 | 0.25 | 0.11 |

Similarly, choice ratios in the control task were not different across sensory modalities, as shown in **Fig. S1**.

#### Model parameters of the value task

**Table S2** is supplementary to **Fig.2F** and illustrates the model parameters *τ*, µ, and σ for each sensory modality in the value task (see also **Figure S2**).

**Table S2.**
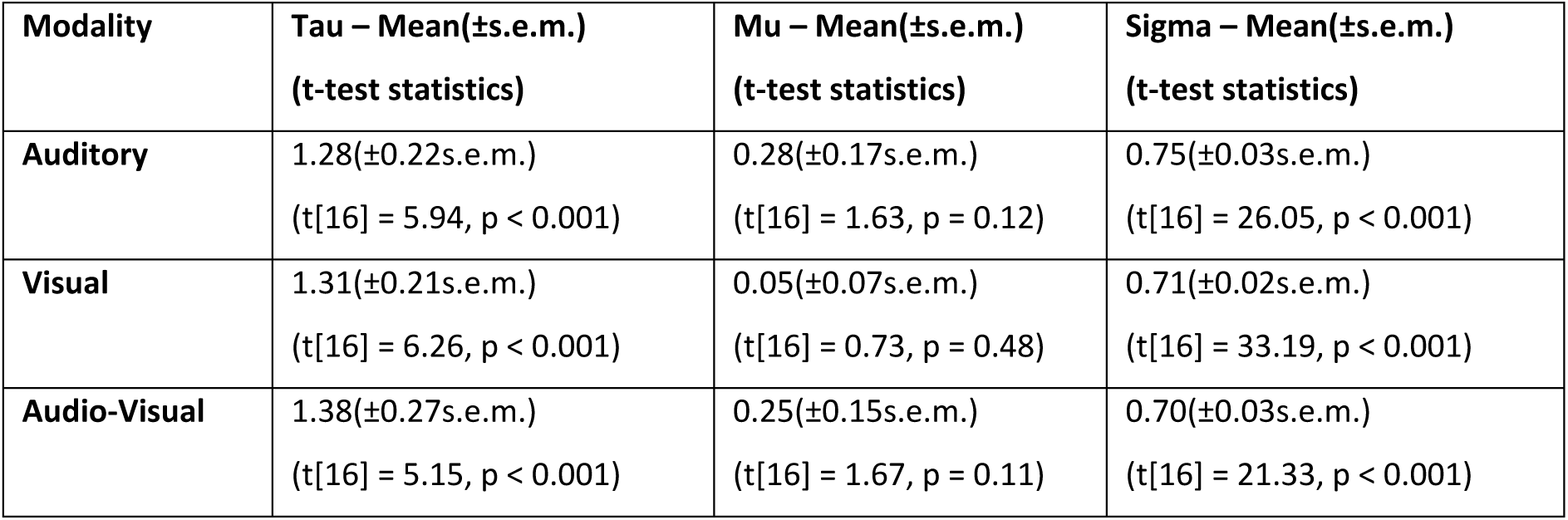
Mean of model-fit parameters across participants and one-sample t-test statistics – for individual conditions (Aud, Vis, AudVis) of the value task.

#### Testing the collinearity of DV and RT

To ensure that the parametric regressors captured dissociable components in pupil response during the value task, we additionally assessed potential collinearity between DV and RT. For each participant, we computed the correlation between trial-by-trial fluctuations in differential value and reaction time across trials. The correlation coefficients across participants were statistically evaluated using a one-sample t-test against zero. No significant correlation was observed at the group level (mean ± s.e.m. = 0.001 ± 0.02, t[16] = 0.03, p = 0.98), indicating that DV and RT captured largely independent sources of trial-by-trial variability within the GLM framework.

#### The effect of value, decision and general task-related regressors on pupil data

**Table S3** is supplementary to **Fig.3F** and shows the pairwise comparisons of different regressor types (DV, RT and EV) and sensory modalities for the pupil data.

**Table S3.**
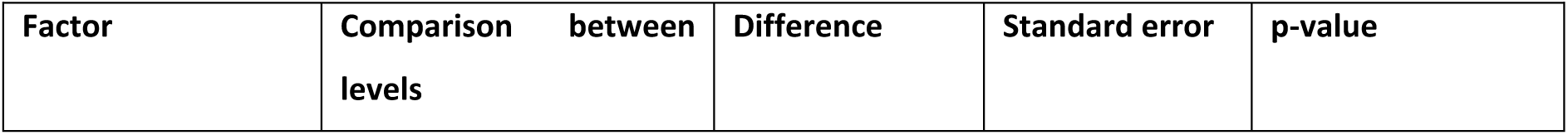
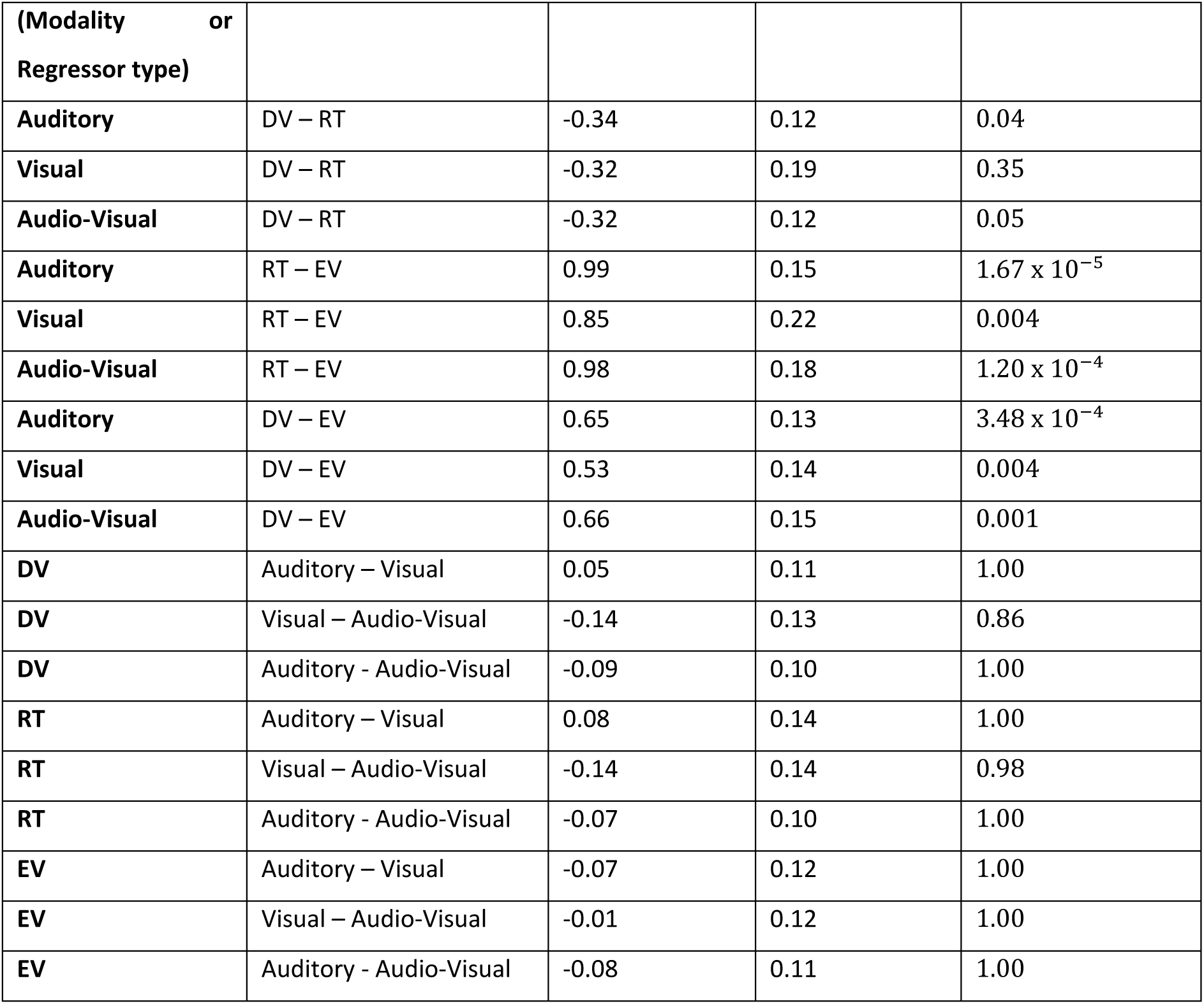
Post-hoc comparisons.

| **Table S3. Post-hoc comparisons** |  |  |  |  |
| --- | --- | --- | --- | --- |
| **Factor** | **Comparison between levels** | **Difference** | **Standard error** | **p-value** |

| (Modality or Regressor type) |  |  |  |  |
| --- | --- | --- | --- | --- |
| **Auditory** | DV – RT | -0.34 | 0.12 | 0.04 |
| **Visual** | DV – RT | -0.32 | 0.19 | 0.35 |
| **Audio-Visual** | DV – RT | -0.32 | 0.12 | 0.05 |
| **Auditory** | RT – EV | 0.99 | 0.15 | $1.67 \times 10^{-5}$ |
| **Visual** | RT – EV | 0.85 | 0.22 | 0.004 |
| **Audio-Visual** | RT – EV | 0.98 | 0.18 | $1.20 \times 10^{-4}$ |
| **Auditory** | DV – EV | 0.65 | 0.13 | $3.48 \times 10^{-4}$ |
| **Visual** | DV – EV | 0.53 | 0.14 | 0.004 |
| **Audio-Visual** | DV – EV | 0.66 | 0.15 | 0.001 |
| **DV** | Auditory – Visual | 0.05 | 0.11 | 1.00 |
| **DV** | Visual – Audio-Visual | -0.14 | 0.13 | 0.86 |
| **DV** | Auditory - Audio-Visual | -0.09 | 0.10 | 1.00 |
| **RT** | Auditory – Visual | 0.08 | 0.14 | 1.00 |
| **RT** | Visual – Audio-Visual | -0.14 | 0.14 | 0.98 |
| **RT** | Auditory - Audio-Visual | -0.07 | 0.10 | 1.00 |
| **EV** | Auditory – Visual | -0.07 | 0.12 | 1.00 |
| **EV** | Visual – Audio-Visual | -0.01 | 0.12 | 1.00 |
| **EV** | Auditory - Audio-Visual | -0.08 | 0.11 | 1.00 |

#### Cross-task comparison of regressor types for the pupil data

**Table S4** is supplementary to the data shown in **Fig. 4** and shows the pairwise post-hoc comparisons of regressor types and tasks (value, control).

**Table S4.** Post-hoc comparisons.

| **Table S4. Post-hoc comparisons** |  |  |  |  |
| --- | --- | --- | --- | --- |
| **Factor**<br>**(Task or Regressor type)** | **Comparison between levels** | **Difference** | **Standard error** | **p-value** |
| **Value Task** | DV – RT | -0.32 | 0.10 | 0.01 |
| **Value Task** | RT – EV | 0.89 | 0.13 | $1.15 \times 10^{-5}$ |
| **Value Task** | DV – EV | 0.57 | 0.09 | $2.74 \times 10^{-5}$ |
| **Control Task** | DV – RT | -0.20 | 0.13 | 0.38 |
| **Control Task** | RT – EV | 0.32 | 0.24 | 0.62 |
| **Control Task** | DV – EV | 0.12 | 0.14 | 1.00 |
| **DV** | Value – Control | 0.16 | 0.07 | 0.04 |
| RT | Value – Control | 0.28 | 0.11 | 0.02 |
| EV | Value – Control | -0.29 | 0.13 | 0.03 |

## Supplementary Figures

**Figure S1.**
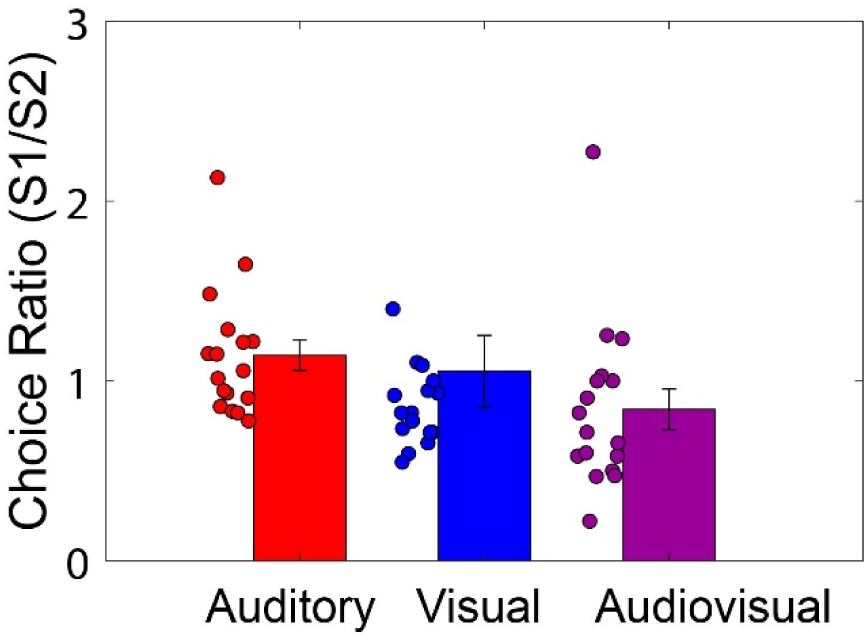
Choice behavior in the control task across sensory modalities. Mean choice ratio (*S*_1_/*S*_2_) across participants for the control task, shown separately for auditory (red), visual (blue), and audio–visual (purple) conditions under an equal switch ratio (1:1). Bars represent group means, and dots indicate individual participant data. Error bars denote ± s.e.m. across participants. Choice behavior was evenly distributed across modalities, consistent with the instruction-driven nature of the task and the absence of value-based learning.

**Figure S2.**
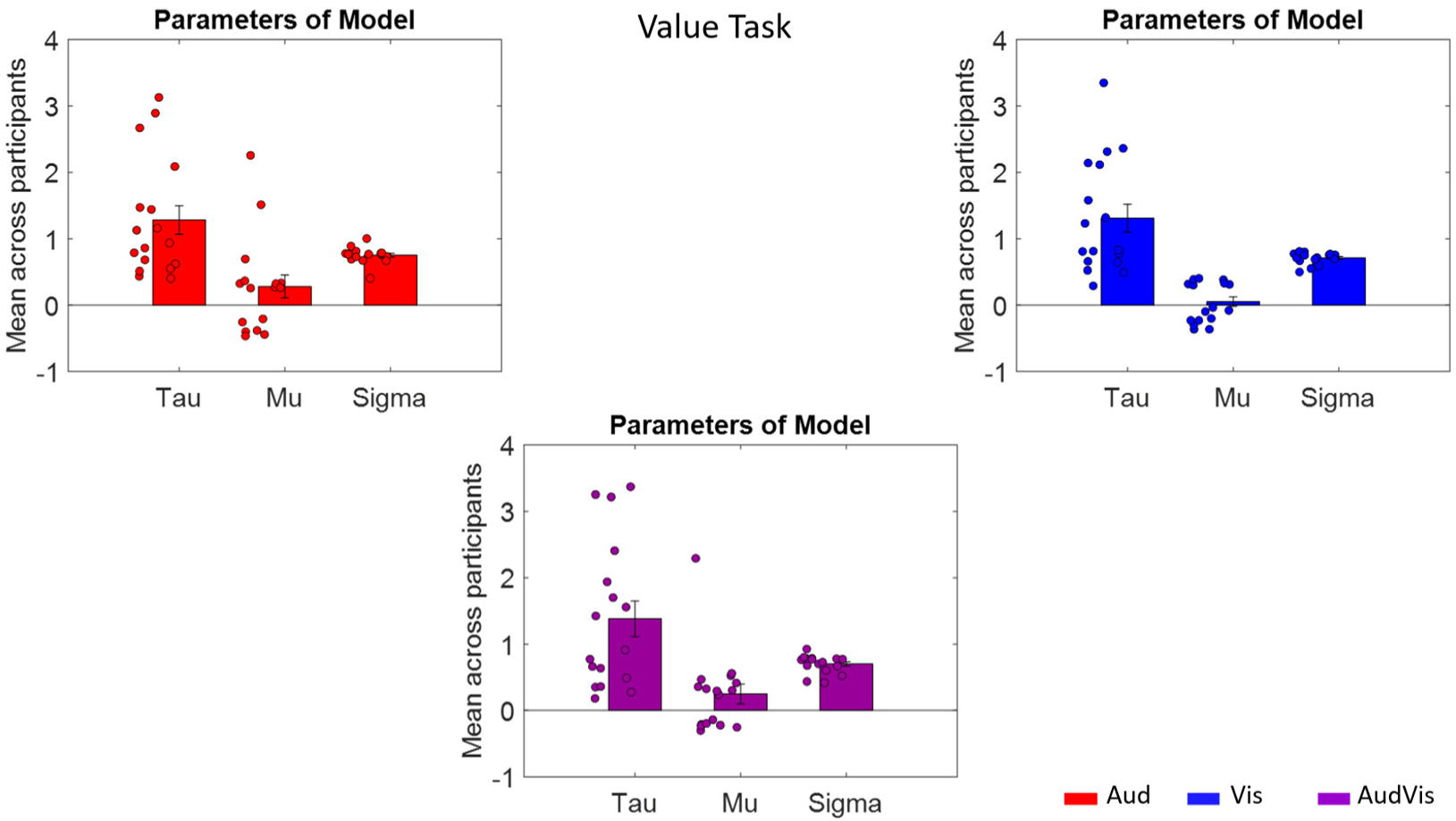
Model parameters across sensory modalities in the value task. Mean estimates of model parameters—time constant (*τ*), bias (μ), and sensitivity (σ)—across participants for auditory (red), visual (blue), and audio–visual (purple) conditions in the value task. Each panel corresponds to one sensory modality, with bars representing group means and dots indicating individual participant estimates. Error bars denote ± s.e.m. across participants.

